# Involvement in family planning service utilization and associated factors among married men at Debre Tabor Town Northwest, Ethiopia, 2017

**DOI:** 10.1101/863712

**Authors:** Tadesse Wuletaw Demissie, Arya Mesfin Nigatu, Enatinesh Mesfin Tegegne

## Abstract

**Background:** Men’s involvement in family planning can be either as a user of male contraceptive methods and or supportive partner of female users.

It is very important to improve women’s health in particular and reproductive health in general. In some developing countries, levels of communication on this issue are low for decision-making. Communication and discussion positively affect contraceptive uptake and continued use, whereas failure to communicate limits couples’ effective and sustained contraceptive use. Unmet need for FP suggested that unwanted pregnancy and unsafe abortion are the main causes of maternal mortality in Ethiopia.

**Objective:** To assess men’s involvement in family planning service and associated factors among married men at Debre Tabor Town, Northwest Ethiopia.

**Methods:** A community based cross-sectional study design was conducted from April to June. 2017 at Debre Tabor Town. Simple random sampling method was recruited then systematic sampling technique was used to include 382 married males. Data were collected by face to face interview using a structured questionnaire. The data were entered into epi. Info 7 and descriptive statistics were analyzed using SPSS version 21 statistical software package.

**Result:** From three hundred eighty-two participants, 373 participants with 97.6% response rate participated in the study. The age range of the participant was from20 to 65years, the mean age was 38.6 with the standard deviation of 7.8. Majority of the participants were orthodox 358(96%). One hundred twenty-four (33.2%) were greater than secondary education and from the occupation, farmers took the least rank 4 (2.4%). The magnitude of male involvement in family planning was 254(68.1%), 370 (99 .2%) of the participants had information on different family planning methods. Adjusting all other factors for the last model, educational status AOR [95%CI] = 2.39 [1.084, 5.260], source of information information AOR [95%CI] =1.88 [1.016, 3.485], men’s approval AOR [95%CI] = 0.07 [0.036, 0.134], ever used contraceptive AOR [95%CI] = 0.21[0.064, 0.705] were found to be associated with men’s involvement.

**Conclusion:** The level of male involvement was moderate, but their actual utilization is low.

## Introduction

### Background

Men’s involvement in reproductive health has two major sides, as men give sufficient support in needs, choices and rights to their partners in reproductive health and fertility control, on the other hand, men’s owns reproductive health issues related to knowledge, contraceptive use and safe sexual behaviours [1].

Family planning has got acknowledgement in the 1990s by many women’s health programs and must be viewed in the broader context of reproductive health; the International Conference on Population and Development (ICPD) held in Cairo 1994 noted that special efforts ought to be made to emphasize men’s shared responsibility and promote their active involvement in responsible parenthood, sexual and reproductive behaviour, including family planning[2].

Men’s participation in Family planning includes, encourage and support their partners (wives) in contraception and encourage peers to use family planning and influence the policy environment to be more conducive for male-related programs[3].

Available studies showed that in many developing countries males often dominate in making important decisions in the family, which include those concerned with reproduction, family size, and contraceptive use [4].

Spousal communication on contraception and reproductive goals suggests that the couple has an egalitarian relationship. Studies have shown that couples who discuss the number of children they desire or the use of family planning are more likely to use a contraceptive and achieve their reproductive goals than those who do not have communication each other[5].

Men have the final say in decision making about family size and the use of contraceptives[6]. A considerable discordance between spouses on questions of family planning and desired family size has also been identified; in some developing countries, levels of communication on these topics are low[7].

African men are not only head of the household but also are overall responsible for families; men have more influence on reproductive decision since they typically control the family asset[8].

In Ethiopia, FP was initiated four decades ago; however, even after such a long period, the service has been amongst the lowest in Africa with 42% contraceptive prevalence rate (CPR) and 18.8% unmet need for FP with current time[9]. Several factors are incriminated for the low coverage of FP services including the desire to have more children, lack of knowledge about contraceptive use and where to find contraceptives, health concerns, religious prohibition, husband opposition and low involvement of males[10].

## Methods

The community-based cross-sectional study design was employed.

### Settings

The study was conducted at Debre Tabor town. Debre Tabor town administration is one of the 13 Districts and five town administrations found in south Gondar administrative zone, Ethiopia. It is located 105km away from Bahir Dar and 666 km away from Addis Ababa. Debre Tabor town is additionally divided into 4 kebeles. The town has a total population of 83,082 of whom 39,781 are males and 43,301 are females (BOFED 2015).

The total number of households in the town is 13200 and there is 1 governmental Hospital, 3 governmental health centres, 3 private clinics and 2 private pharmacies.

The study was conducted from April to June.30/2017 in Debre Tabor Town administration of South Gondar, Ethiopia.

### Study population and sampling

All married men who are living in Debre Tabor town administration.

The sample size was calculated using a single population proportion formula using an estimated 65.5% proportion of male involvement [11] with a 95% confidence level, 5% marginal error and considering 10% none response rate, then the final sample was 382.

Debre Tabor town has four kebeles. Currently, there are about 13200 households in the town with 4624 married males. Simple random sampling (lottery method) technique was undertaken to select households. The four kebeles were included in the study. Participants were allocated proportionally to get 382 married males from 4624 married males in the four kebeles. Married males were selected systematically every 12 households in each kebele.

### Operational definition

- **Current use of contraceptive**: Those respondents who are using the contraceptive method or whose partners are using the contraceptive method during the data collection period.
- **Men involvement**: - Spousal communication, approval, support of the husband and both partners usage of contraceptive and score above the mean.
- **Married man**: - A man who lived together with a woman with a legal relationship for greater than 6 months.
- **Good men’s attitude towards family planning**:-respondents who answered agree and above on the attitude questions

### Data collection procedures and instruments

Data were collected using a structured questionnaire with face to face interview technique

### Data management and analysis

The data were entered using Epi. Info version 7; SPSS version 21 was being used for data cleaning and analysis.

### Ethical considerations

Ethical clearance was obtained from the ethical review committee of DTU. The participants were informed orally the informed verbal consent was obtained from each participant.

## Result

### Socio-demographic characteristics

From 382 participants invited to participants, 373 participants were participated in the study yielding 97.6%. The age of the participants ranged from 20-65 years, the mean age was 38.6 with the standard deviation of 7.8. Majority of the participants were orthodox 359 (96.2%) and 11 (2.2%) were Muslims. With respect to educational status most participants 124 (33.2%) were greater than secondary education, 11 (2.9%) of the participants were unable to read and write and with regard to occupation 163 (43.7%) were governmental employees, 22 (5.9%) of them were unemployed and farmers took the least rank 9(2.4%). The highest frequency of age at marriage is 30 years 130 (34.9%) and most of the respondents 122(32.7%) have 3-4 children and the least 41(11%) of them have more than five children (See: Table: 1).

**TABLE-1:**
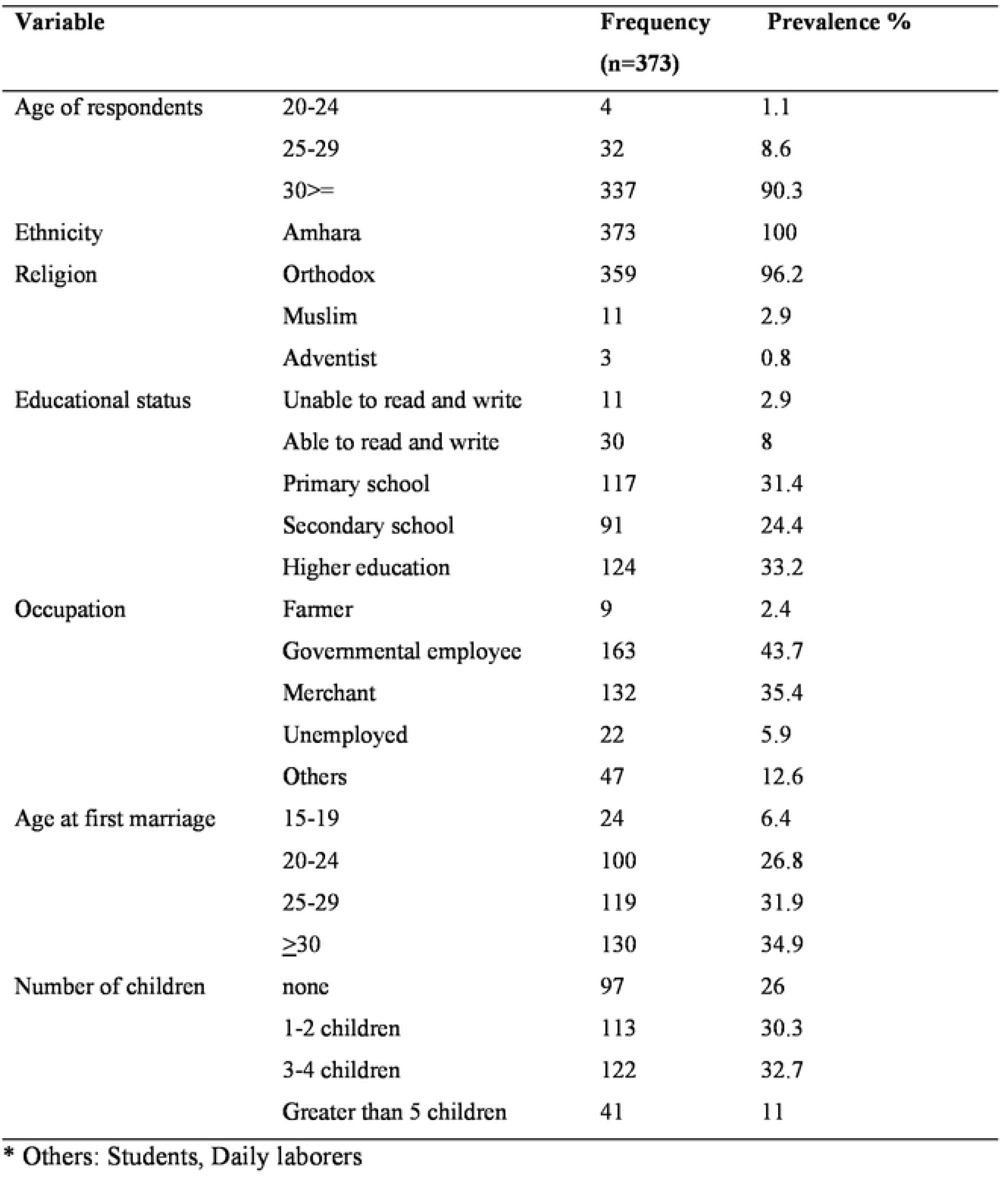
FREQUENCY DISTRIBUTION OF SOCIO DEMOGRAPHIC CHARACTERISTICS AMONG MARRIED MEN AT DEBRE TABOR TOWN, NORTHWEST ETHIOPIA, 2017.

Married men had information on family planning methods by different Media for them and their spouses; 99.2 % of the participants had information on different family planning methods; Parents took the first rank to be a source of information and television accounts least (See:Table:2).

**TABLE-2:**
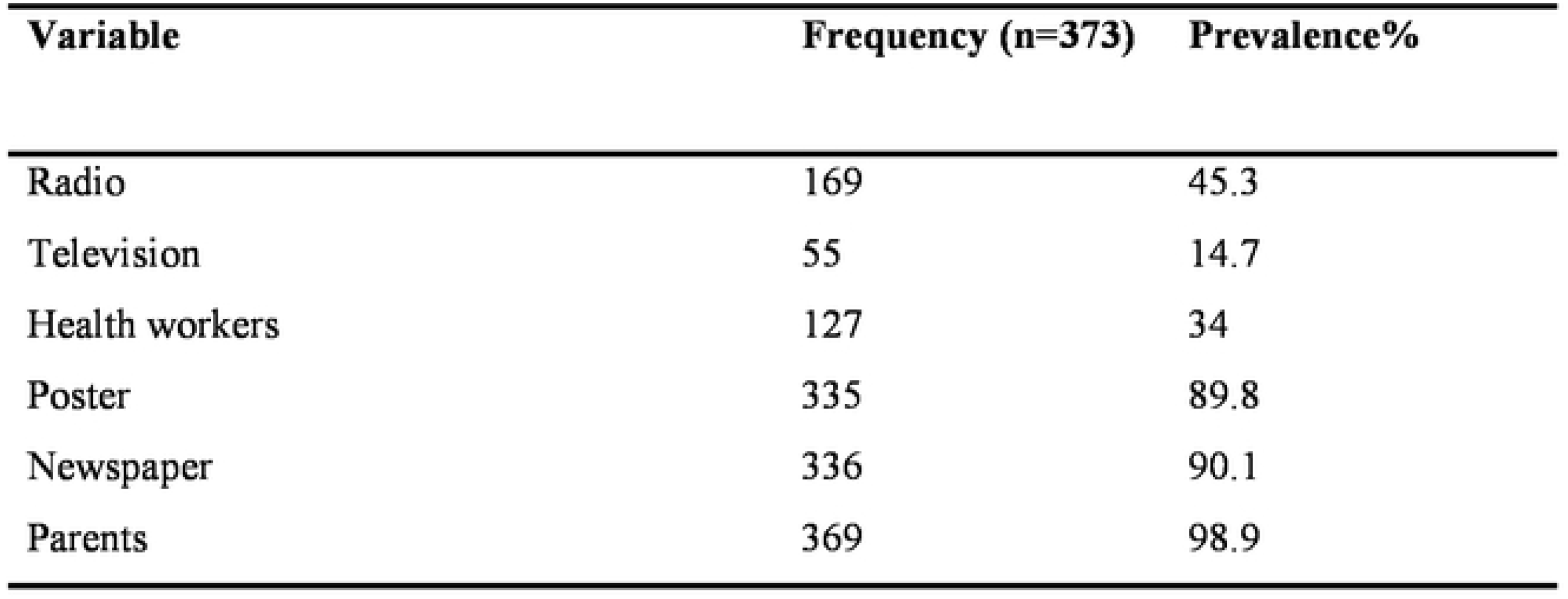
FREQUENCY DISTRIBUTION OF SOURCES OF INFORMATION ABOUT FAMILY PLANNING METHODS AMONG MARRIED MEN AT DEBRE TABOR TOWN, NORTHWEST ETHIOPIA,2017.

| **Variable** | **Frequency (n=373)** | **Prevalence%** |
| --- | --- | --- |
| Radio | 169 | 45.3 |
| Television | 55 | 14.7 |
| Health workers | 127 | 34 |
| Poster | 335 | 89.8 |
| Newspaper | 336 | 90.1 |
| Parents | 369 | 98.9 |

**TABLE-3: BIVARIATE AND MULTIVARIATE ANALYSIS FOR FACTORS ASSOCIATED WITH MEN INVOLVEMENT IN FAMILY PLANNING METHODS AMONG MARRIED MEN AT DEBRE TABOR TOWN, NORTHWEST ETHIOPIA, 2017.**

### Men’s attitude and or support towards contraceptives

Among the study participants, 358 (96%) of them had a good attitude or positive support on contraceptive utilization for them and their partners.

### Men’s involvement in family planning

Men’s involvement on family planning was 254(68.1%) 95%CI= [63.4%, 73.2%]. Most of the men had awareness on F/P, support their spouse, 93.8% of them discuss about F/p and 71.8% approve it, but only 28(7.5%) of the men had used family planning; particularly 16 (4.3%) were used condom which is lower than the national prevalence in 2016 EDHS (11%) and 12(3.2%) of them used abstinence and none of them was used male sterilization. Sixty-eight per cent of female partner took contraceptives which is higher than the national CPR (36% in 2016 EDHS); some of the reasons mentioned not to take contraceptives were: fear of side effects, partner opposition, lack of awareness and desire for more children (see: Fig-1).

**Fig.1:**
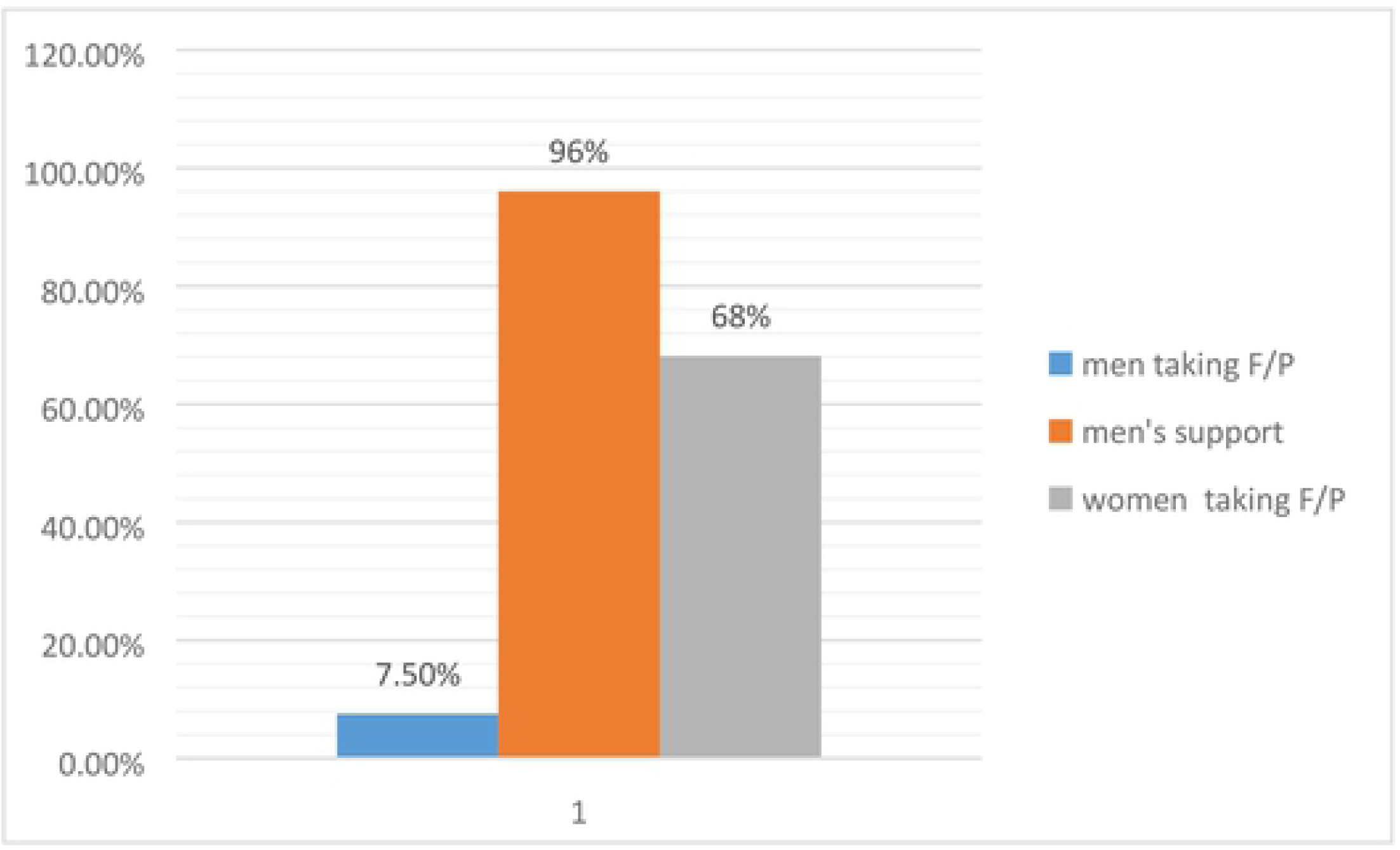
Men’s involvement and support on family planning.

Even though married men had information, awareness and supported on family planning for them and their spouses, about 254(68.1%) of men were involved in family planning (see: Fig-2).

**Fig.2:**
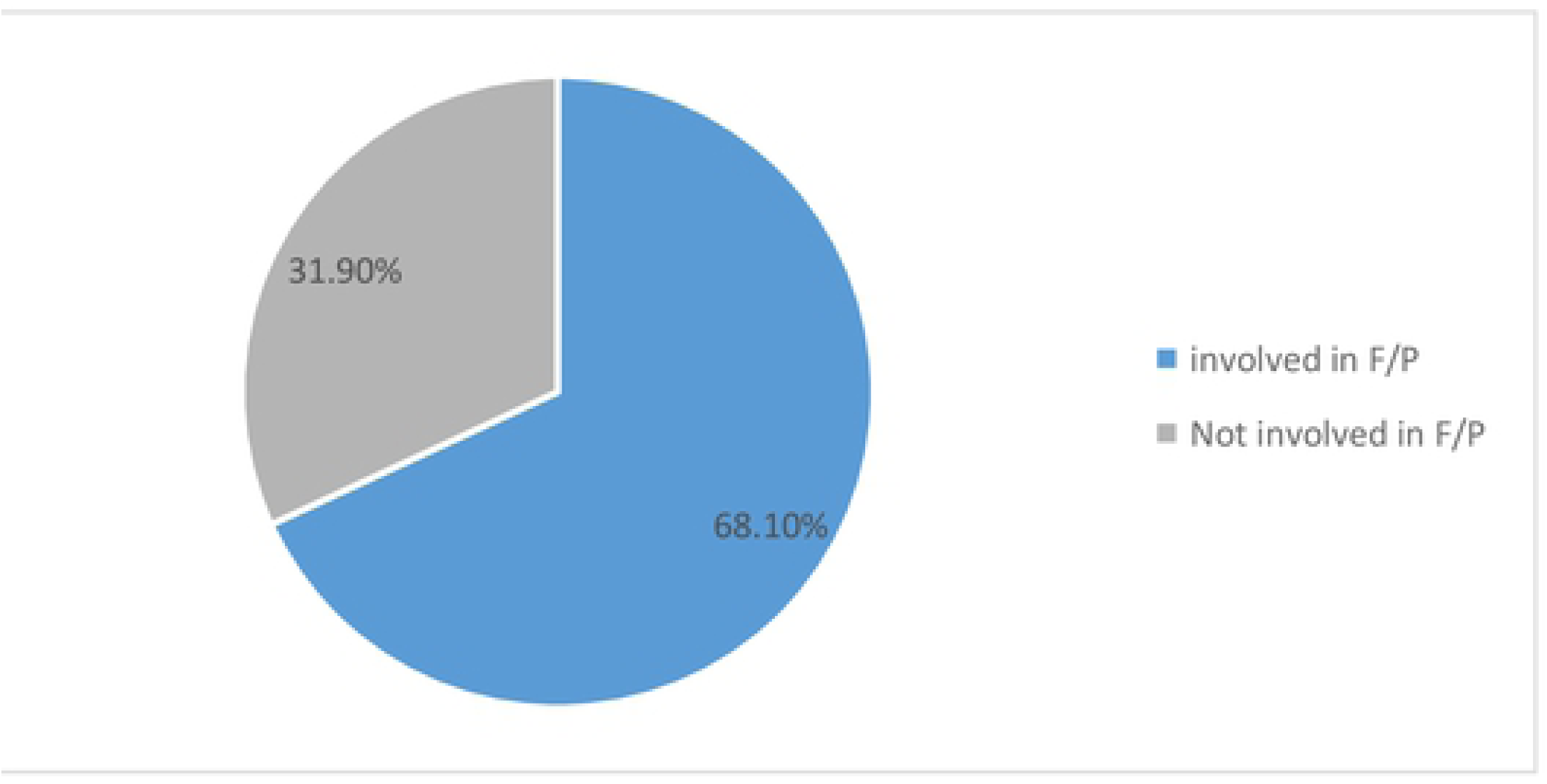
Men’s involvement in family planning.

### Factors associated with men’s involvement in family planning

Bivariate analysis was used to explore the association between men’s involvement in family planning with each of the determinants factors. Educational status of married men, sources of information’ about F/P, know male type of contraceptive method, ever informed their partner to use F/P, had desire to know more about F/P methods, men’s approved for family planning method, men’s ever usage of F/P method and previous usage of F/P methods by female partner were met the minimum criteria (P<0.2) for further multivariate logistic analysis.

Factors which had bivariate associations at p-value < 0.2 were then entered into multivariate logistic regression analysis. The variables associated by multivariate logistics analysis (p ≤ 0.05) with men’s involvement in family planning were: Educational status of married men, sources of information’ about F/P, married men ever informed their partner to use F/P, had desire to know more about F/P methods, men’s approved for family planning method, ever used of F/P method by male partner and previously usage of F/P methods by female partner (see: Table 3).

**TABLE-3:**
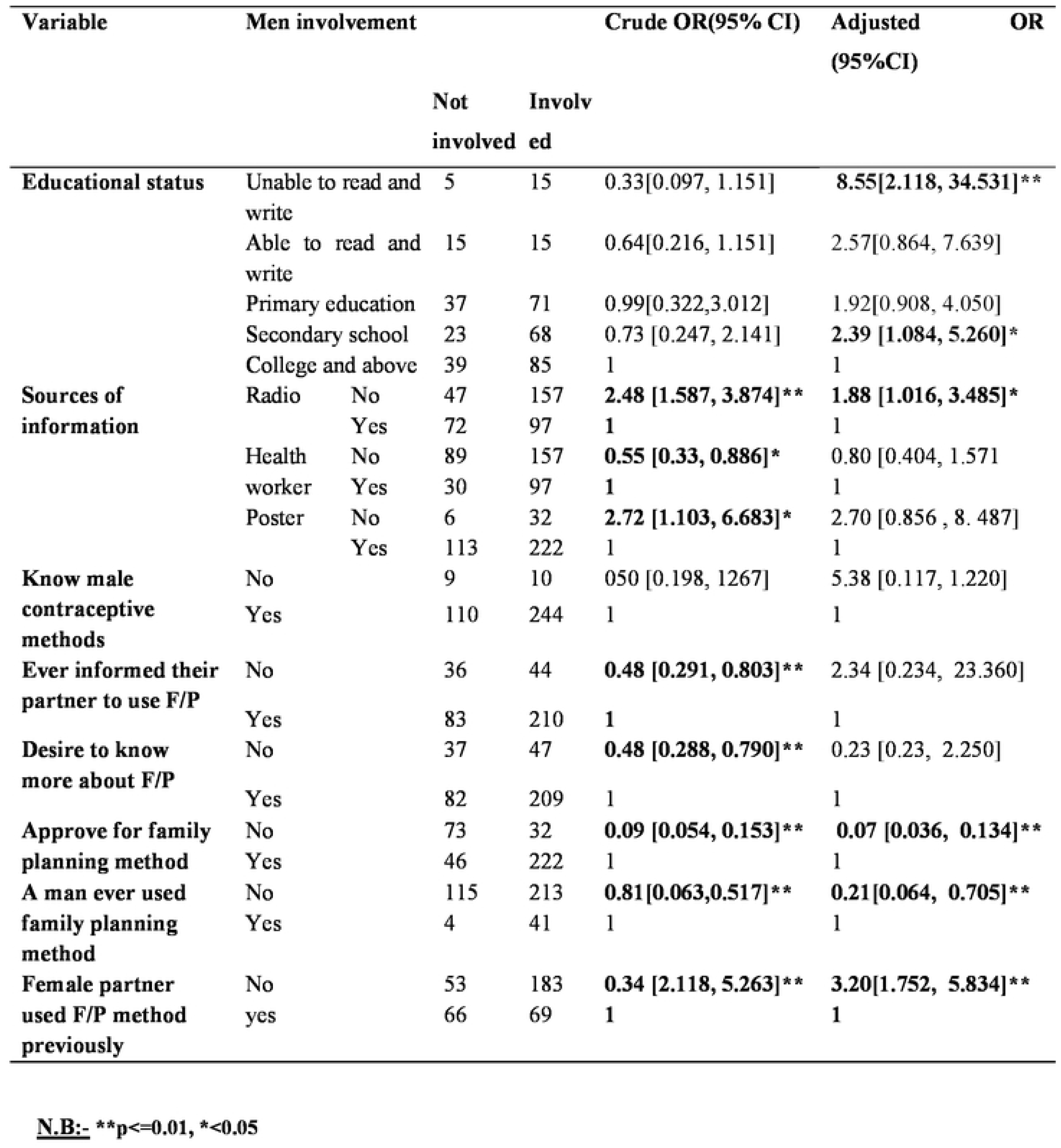
BIVARIATE AND MULTIVARIATE ANALYSIS FOR FACTORS ASSOCIATED WITH MEN INVOLVEMENT IN FAMILY PLANNING METHODS AMONG MARRIED MEN AT DEBRE TABOR TOWN, NORTHWEST ETHIOPIA, 2017.

## Discussion

### The magnitude of men’s involvement in family planning service among married men

A community-based cross-sectional study was conducted to assess the magnitude of married men’s involvement in family planning and associated factors at Debre Tabor Town. In this study, the magnitude of married men’s involvement in family planning was 68.1% which is nearly in line with the study done in Bangladesh (63.2)[12] and Vietnam(63.7%)[13].

The current study was higher than the study done in West Pokot, Kenya (52%) [14] turkey (30%)[15], only 4.8% were involved in Nigeria[16], in west Shewa zone, Ethiopia(36%)[17] and Afar, Ethiopia [18]. This could be due to governmental concern, economical factor, individual motivation and discussion and approval of the service by male partners.

The result is lower than the study conducted in rural Vietnam (74.4%) [19]. This could be due to less shared responsibility, educational status and poor attitude towards male participation in family planning service.

Participants claimed that they have used family planning were only 7.5% which is lower than the study conducted in Debre Markos town, Ethiopia (8.4%) in eastern Tigray Ethiopia (15%)of the participants were using family planning methods directly[20, 21], and in South-Eastern Turkey(39.6%)[22], but higher than the study done in Southern Ethiopia (5%) [11],in west Pokot Kenya (6%)[14]. This is due to awareness on family planning and accesses to family methods, economical biological and socio-cultural impacts of men furthermore; governmental and couple concerns on family planning methods.

Based on access to information on family planning about 99.2 % married males had got information on family planning methods via different Media which is in line with the study done in Debre Markos, Ethiopia (99.2%) [20]but higher than the study done in Malegedo Town, Oromia, Ethiopia(89%)[17]. This could be the access to mass media.

The study result is higher than the study conducted in India (98% [23], in Southern Turkish 43.2% [22], in Ghana(89%)[24],in North Shewa, Ethiopia (93.9%) [25] in Woliyta Sodo Southern Ethiopia((96%)[11].in Tigray, Northern-Ethiopia 62.9% of the participants were informed about F/p[21, 24]. More involvement of health care providers to motivate, governmental concerns and media motivation to create awareness furthermore individual motivation could be mentioned.

### Factors associated with men’s involvement in family planning

According to this study the multivariate analysis of logistic regression for men’s involvement in family planning pointed out that; Educational status of married men, sources of information’ about F/P, men ever informed their partner to use F/P, had desire to know more about F/P methods, men’s approved for family planning method, men’s ever usage F/P method and female partner used F/P methods previously were found to be significantly associated with men involvement on family planning.

The study result revealed that educational status of male partner was found to be one factor for men’s involvement in family planning; those respondents who were unable to read and write AOR [95%CI] = 8.55[2.118, 34.531] and those in secondary education level AOR [95%CI] = 2.39 [1.084, 5.260] were 8.6 and 2.3 times more likely to be involved in family planning service respectively as compared to higher education level. This result is consistent with the study done in Eastern Tigray, Ethiopia [21], in kenya[14] and Turkey[22]. This is because individual concern could not be affected by education, moreover, this could be the low economic status of the individuals.

Source of information is also one significant factor. Radio as a source of information AOR [95%CI] =1.88 [1.016, 3.485] was 1.8 times more likely affected men to be involved in family planning than those did not have information. This result is consistent with the study done in Malegedo Town, Oromia, Ethiopia[17].

In this study, men’s approval for family planning was strongly associated with men’s involvement in family planning. Those men approve family planning for their partner AOR [95%CI] = 0.07 [0.036, 0.134] were by 93% less likely to involve in family planning than those not approve family planning. This result is inconsistent with the study conducted in Bench Maji Zone, Ethiopia and Debre Markos, Ethiopia [26, 27]. This may be due to the accessibility of information and shared responsibility; female partners were taking responsibility for family planning.

Ever used family planning method was found to be a significant factor. Those male partners ever used family planning AOR [95%CI] = 0.21[0.064, 0.705] were by 79% less likely for the involvement in family planning than those did not use family planning This could be due to the need for more children, less concern for discussion and disapproval of partner contraceptive usage.

Previously usage of F/P method was strongly associated with men’s involvement in family planning. Female partner used F/P method previously AOR [95%CI] = 3.20[1.752, 5.834] were 3.3 times more likely to be involved in family planning. This may be the experience sharing for male partners and shared responsibility.

### Limitation of the study

As the study was cross-sectional, we cannot infer the temporal relationship between information and family planning involvement of male partners.

Men directly involved in family planning with only abstinence and condom this may need further research mainly qualitative research.

## Conclusion and recommendation

In this study, the level of male involvement was moderate, but their actual utilization is low. Inaccessibility to the services, the desire to have more children, socio-economical, biological and technological factors were found to be the reasons for the male involvement in family planning services utilization.

Governmental and non-governmental organizations, service providers, policymakers and relevant stakeholders should ensure availability, accessibility and sustained advocacy for use of family planning services. The family planning programs should incorporate males in the uptake of family planning services.

## Acronyms & abbreviations

BOFED: Bureau of finance and economic development
BSc: Bachelor of science
CI: Confidence interval
CPR: Contraceptive prevalence rate
DTU: Debre Tabor University
EDHS: Ethiopian demographic and health survey
FP: Family planning
FIG: Figure
ICPD: International conference on population development
OR: Odds ratio
p-value: Probability value
SPSS: statistical package for social science
SSA: Sub-Saharan Africa
STI: Sexually Transmitted Infections
WHO: World health organization

## Authors’ contributions

**TW** designed the study, participated in the data quality control, performed analysis and interpretation of draft of the paper and prepared the manuscript. **AM** assisted the design, approved the proposal, and revised drafts of the paper. **EM** also assisted the design, approved the proposal, and revised drafts of the paper. All authors contributed to and approved the final manuscript.

## Ethical approval and consent to participants

Ethical clearance was obtained from Debre Tabor University Ethical Review Board, Permissions was being taken from the South Gondar Zonal Health Department. The necessary explanation about the purpose of the study and its procedure was given to the participants and then, informed verbal consent was obtained from the respondents. The study participants had got information about that, they have the full right not to participate in the study if they are not willing. Anonymity was explained clearly for the participant to ensure confidentiality.

## Data availability

The data set and other resources will be available online as per the rule and regulation of the journal.

## Acknowledgement

We would like to acknowledge Debre Tabor University, Department of Nursing for fulfilling all things to go through and facilitating this study.

## Reference

1. J C, K Y, R R. Men’s involvement in family planning in rural Bangladesh. 2008.

2. Butto1 D, Mburu S. Factors Associated with Male Involvement in Family Planning in West Pokot County, Kenya. Universal Journal of Public Health 2015;3(4):160–8.

3. Kamal MM, Islam1 MS, Alam MS, Hassan ABME. Determinants of Male Involvement in Family Planning and Reproductive Health in Bangladesh. American Journal of Human Ecology. 2013;2:83–93.

4. Commission NP. population census of the Federal Republic of Nigeria, Abuja, Nigeria. 1998.

5. Chem Ga. male involvement in family planning 2003.

6. Puri S, Walia D, Mangat, Kalia Mea. Current Scenario of Contraception and Indian men. Middle East Journal of Nursing. 2010 4(1).

7. A.Oyediran K, Ishola GP, J.Feyisetan B. Factor Affecting Ever-Married Men’s Contraceptive Knowledge and Use in Nigeria Journal of Biosocial Science. 2002;34:497–510.

8. Amon I. THE PARTICIPATION OF MARRIED MEN IN DECISION MAKING ON HOUSEHOLDS’ FAMILY PLANNING METHODS IN MBEYA CITY. 2015.

9. W A, A A, D D. The involvement men an application of the transtheoretical model in family planning in Wolayeta Soddo town south Ethiopia. Asian J Med Sci. 2010;2(2):44–50.

10. Cp G, Si C, Belhadj-El, H G. Male Involvement in Reproductive Health, Including Family Planning and Sexual Health. 1995.

11. 1W. Abraham AAaDD. The Involvement of Men in Family Planning An Application of Transtheoretical Model in Wolaita Soddo Town South Ethiopia. Asian Journal of Medical Sciences 2010.

12. Mohammad Mostufa Kamal1* MSI, Muhammad Shafiul Alam2and A. B. M Enamol Hassan3 Determinants of Male Involvement in Family Planning and Reproductive Health in Bangladesh American Journal of Human Ecology 2013;2.

13. Ha BT, Jayasuriya R, Owen N. Male involvement in family planning in rural Vietnam: an application of the Transtheoretical Model. Health Educ Res. 2003;18(2):171–80.

14. Butto D, Mburu S. Factors Associated with Male Involvement in Family Planning in West Pokot County, Kenya. Universal Journal of Public Health. 2015;3(4):160–8.

15. Sevkat Bahar BGaAA. Male involvement in family planning in Turkey. 1998;19.

16. Ademola Adelekan PO, 2 and Elizabeth Edoni3. Male Involvement in Family Planning: Challenges and Way Forward. Hindawi, International journal of population research. 2014;2014.

17. Demissie DB, Kurke ATAAA, Awel A, Oljira K. Male Involvement in Family Planning and Associated Factors among Marriedin Malegedo Town West Shoa Zone, Oromia, Ethiopia. planning. 2016;15.

18. Mebrahtu Kalayu Chekole1 ZHK, Araya Abrha Medhanyie2, Mussie Alemayehu Gebreslassie2 and Afework Mulugeta Bezabh2. Husbands’ involvement in family planning use and its associated factors in pastoralist communities of Afar, Ethiopia Reproductive health. 2019.

19. Ha BT, Jayasuriya R, Owen N. Increasing male involvement in family planning decision making: trial of a social-cognitive intervention in rural Vietnam. Health Educ Res. 2005;20(5):548–56.

20. Mhretie kassa AAaMG. Level of male involvement and associated factors in family planning service utilization at Debre Markos Town Northwest Ethiopia BMC international health and Human Right. 2014;14.

21. Bayray A. Assessment of male involvement in family planning use among men in the southeastern zone of Tigray, Ethiopia. Scholarly Journal of Medicine. 2012;2(2):1–10.

22. Kisa S, ZeyneloĞLu S, DelIBaS L. Influence of age on the usage of family planning methods by Turkish married men living in southeastern Turkey. Turkish Journal of Medical Sciences. 2013;43:756–63.

23. Char A. Male Involvement in Family Planning and Reproductive Health in Rural Central India. 2011.

24. 1Ghana Health Service Td, Upper East Region, Ghana Factors influencing the uptake of family planning services in the Talensi District, Ghana Panafrican medical journal. 2014.

25. Negash W, Dessalegn M, Yitayew B, Demsie M, Wagnew M, Nyagero J. Reproductive health service utilization and associated factors: the case of north Shewa zone youth, Amhara region, Ethiopia. Pan Afr Med J. 2016;25(Suppl 2):3.

26. Henok A, Takele E. Assessment of Barriers to Reproductive Health Service Utilization among Bench Maji Zone Pastoralist Communities. Ethiop J Health Sci. 2017;27(5):523–30.

27. Kassa M, Abajobir AA, Gedefaw M. Level of male involvement and associated factors in family planning services utilization among married men in Debremarkos town, Northwest Ethiopia. BMC international health and human rights. 2014;14(1):33.

